# Alcohol or stress exposure during late adolescence impairs risk assessment later in life, disrupting the ethological structure of behavior in mice

**DOI:** 10.64898/2026.01.09.698658

**Authors:** Beatriz Deo Sorigotto, Isabel Marian Hartmann de Quadros, Karina Possa Abrahao

## Abstract

**Background:** Decision-making is a key component of adaptive behavior and relies on risk assessment as a critical process that enables organisms to evaluate threats before exploring potentially dangerous areas. During adolescence, this process is particularly vulnerable to disruption by external factors such as stress and substance use, potentially leading to changes in the structure of defensive and exploratory behaviors, potentially leading to riskier choices.

**Methods:** We investigated whether repeated ethanol intoxication or pharmacological stress during late adolescence (7-8 weeks of age) alters overall behavioral structure, focusing on risk assessment behaviors, in adulthood. Male and female C57BL/6 mice received four intermittent injections of saline, ethanol (4.0 g/kg), or the α2-adrenergic antagonist yohimbine (2.0 mg/kg). In early adulthood (9th week), animals were tested in the Light–Dark Box and Elevated Plus Maze. In addition to traditional anxiety-like behavior and exploratory measures, risk assessment behaviors (stretches and head outs) were classified as NoGo (risk evaluation without entering the aversive area) or Go (risk evaluation followed by exploration), and behavioral organization was examined using first-order Markov chain analyses.

**Results:** Both ethanol and yohimbine exposure produced persistent reductions in NoGo risk assessment behaviors in adulthood, in both sexes, with minimal effects on classical anxiety-like measures. Correlation and transition probability analyses revealed a reorganization of behavioral structure, characterized by reduced transitions toward risk assessment events and a bias toward risk-taking actions.

**Conclusions:** These findings indicate that repeated late adolescent alcohol intoxication and stress exposure induce long-lasting alterations in risk assessment and decision-making strategies.

## Introduction

Decision-making is an adaptive process shared by all organisms, involving the evaluation and integration of internal and external information to guide behavior. A central aspect of decision-making is risk assessment, a defensive behavioral process through which an organism, confronted with potential or ambiguous threats, detects and analyzes both the threat stimulus and the surrounding context (Blanchard and Blanchard, 1989; Blanchard et al., 2011). This process integrates features such as the type, location, and proximity of the threat, along with environmental factors like the availability of escape routes or shelter, to guide the selection of the most adaptive defensive strategies, thereby minimizing danger (Blanchard et al., 2011). This process also relies on inhibitory control to suppress impulsive actions and allow more adaptive choices (Loyant et al., 2025). Risk assessment behaviors are considered to be closely associated with anxiety-like behaviors (Carobrez and Bertoglio, 2005) and to reflect a conflict between approach and avoidance tendencies (Blanchard et al., 1991). In rodents, such behaviors are commonly investigated using experimental paradigms that mimic instinctive approach-avoidance conflict, typically through exposures to bright, open areas and elevated platforms. Classical anxiety-related tests like the Light-Dark Box (LDB) and the Elevated Plus Maze (EPM) take advantage of rodents’ innate aversion to brightly lit environments (Bourin and Hascoët, 2003) and open, elevated spaces (Kraeuter et al., 2018). When combined with the intrinsic drive to explore novel environments, these naturally aversive contexts elicit approach-avoidance conflicts and, consequently, the expression of risk assessment behaviors (Thompson et al., 2018; La-Vu et al., 2020).

External factors such as drugs of abuse and stressors can interfere with approach and avoidance behaviors influencing decision-making. For example, the anxiolytic effect of acute ethanol (increased approach and decreased avoidance) has been extensively demonstrated in mice (LaBuda and Fuchs, 2001; Gulick and Gould, 2009) and rats (Bertoglio and Carobrez, 2002; Acevedo et al., 2014) when tested in paradigms such as the EPM. Indeed, ethanol can increase impulsive behaviors in mice (Starski et al., 2019) and impair inhibitory control and decision-making in humans (Caswell et al., 2013; Weafer and Fillmore, 2016). Thereby, it is possible that alcohol can also disrupt risk assessment by shifting the balance between exploratory and avoidance states, leading to maladaptive changes in the ethological structure of behavior. Importantly, these effects are not restricted to acute ethanol intoxication. Chronic binge alcohol drinking followed by protracted abstinence induces persistent behavioral alterations, including reduced avoidance and risk assessment (reflected by increased exploration in aversive areas of the EPM and open field), as well as increased compulsive-like behavior in mice (Rivera-Irizarry et al., 2023). Notably, ethanol exposure during adolescence leads to behavioral alterations that persist into adulthood, including changes in exploratory and emotional behaviors, as well as increased alcohol consumption (Amodeo et al., 2018).

In addition to its direct pharmacological effects, ethanol activates the hypothalamic-pituitary-adrenal (HPA) axis (Quadros et al., 2016), acting as a stressor. Likewise, stressful experiences are powerful modulators of risk assessment and inhibitory control, although their behavioral consequences depend on the nature, intensity, and timing of the stressor. For example, predator odor stress increases anxiety-like behaviors while reducing risk assessment and exploratory activity in rats in the EPM (Adamec et al., 2001), whereas social defeat stress increases risk assessment in rats exposed to the same test (Stickling and Rosenkranz, 2025). Conversely, in humans, social stressors such as financial strain also modulate inhibitory control (Hughes et al., 2024), possibly modulating risk taking behaviors.

These findings suggest that alcohol intoxication and stress exposure can alter how animals evaluate and respond to potentially threatening environments, but they do not rule out the possibility of long-term changes that emerge later in life. This is of particular interest when considering developmental stages, especially adolescence. Neural systems involved in inhibitory control, emotional processing, and stress responsivity undergo protracted maturation throughout adolescence, rendering this period especially vulnerable to environmental perturbations (Spear, 2013; 2018). In fact, adolescence represents a particularly vulnerable developmental period marked by profound behavioral and neurobiological changes and the refinement of adaptive behaviors critical for survival (Laviola et al., 2003). In mice, this period spans approximately from 3 to 8 weeks of age (Bell, 2018), with early adolescence (∼3-5 weeks) corresponding to rapid limbic and motivational circuit maturation, and late adolescence (∼6-8 weeks) aligning with continued development of prefrontal and corticostriatal systems implicated in cognitive control and decision-making (Tarazi et al., 1998; Kasanetz and Mansoni, 2009; Gutierrez-Castellanos et al., 2024; Pöpplau et al., 2024). Notably, these stages are considered broadly analogous to early-mid and late adolescence in humans (Semple et al., 2013; Dutta and Sengupta, 2016).

Exposure to alcohol or stress during adolescence has been associated with long-lasting alterations in affective and motivational behaviors that persist into adulthood. For example, studies in mice show that stress exposure during adolescence increases anxiety-like behaviors later in life (Caruso et al., 2018), and ethanol exposure during early adolescence alters ethanol sensitivity and heightens anxiety-like and risky behaviors in adulthood (Lee et al., 2017; Pichinin et al., 2025, preprint). However, while these studies primarily focus on behavioral outcomes or affective states, less is known about how adolescent alcohol or stress exposure reshapes the organization of decision-making processes themselves, particularly during risk assessment, when mice reach adulthood.

Beyond the quantification of individual behaviors, risk assessment can be better understood integrating it to the decision to take or not take the risk and through the temporal organization and sequencing with other actions. In ethological and ethoexperimental contexts (Blanchard et al., 1989), the way behaviors are chained over time provides critical insight into underlying decision strategies, revealing patterns that are not captured by measures of frequency or duration alone (Brown and de Bivort, 2018). Analytical approaches based on behavioral transitions, which can be represented by Markov chains (Tejada et al., 2010; Sanabria et al., 2019), offer a powerful framework to characterize probabilistic relationships between successive behaviors and to examine how animals navigate approach-avoidance conflicts. By modeling behavior as a sequence of state transitions, these methods have been increasingly used to identify alterations in behavioral organization associated with anxiety, stress, and maladaptive decision-making.

In the present study, we investigated whether repeated heavy ethanol intoxication or pharmacological stress exposure during late adolescence produces persistent alterations in anxiety-like, exploratory and risk assessment behaviors in adulthood. Female and male mice were exposed to ethanol or yohimbine during late adolescence and later evaluated in the LDB and EPM, two paradigms that allow the characterization of approach-avoidance conflict and risk assessment. In parallel, blood ethanol concentrations and plasma corticosterone levels were measured to assess intoxication levels and HPA axis engagement during the exposure period. By focusing on behavioral outcomes that persist beyond the exposure window, this study aims to clarify how adolescent alcohol and stress experiences bias adult decision-making.

## Materials and methods

### Animals

All procedures followed institutional guidelines aligned with national laws and regulations (Comissão de Ética no Uso de Animais; CEUA #7826300522; Universidade Federal de São Paulo, UNIFESP), and were performed in agreement with the NIH Guide for the Care and Use of Laboratory Animals (NIH, 2011). We used adolescent C57BL/6 female and male mice (6 weeks old on arrival; n = 59). All mice were purchased from the Centro de Desenvolvimento de Modelos Experimentais para Medicina e Biologia (CEDEME/UNIFESP, Brazil). Animals were housed in groups of 3-4, separated by sex, in polypropylene cages (44 x 34 x 16 cm) placed in ventilated racks (Alesco, Monte Mor, SP, Brazil) under a 12 h light/dark cycle with lights on at 7 am. Chow (Nuvilab CR-1, Nuvilab, Curitiba, PR, Brazil) and water were available *ad libitum* throughout the study. Prior to the beginning of experiments, animals were allowed to acclimate to the *vivarium* for one week and underwent handling and weighing by the experimenter. Card paper rolls were used for environmental enrichment, and cages were changed once a week.

### In vivo drug exposure protocol

Mice underwent four administrations of saline (Ctrl; 0.9% w/v; Samtec, Ribeirao Preto, SP, Brazil), ethanol (EtOH; 4.0 g/kg, i.p., 20% v/v in 0.9% saline; from 95% ethanol, 01A1082.01.BJ, Labsynth, Diadema, SP, Brazil), or yohimbine (Yoh; 2.0 mg/kg, i.p., 0.08 mg/mL in 0.9% saline; from ≥98% yohimbine hydrochloride, Y3125, Sigma-Aldrich, Munich, BY, Germany), an α2-adrenergic receptor antagonist which increases noradrenaline release and activates the HPA axis (Langer, 1980; Tanaka et al., 2000), over the course of two weeks (during the 7th and 8th weeks of age), with a minimum interval of 48 h between injections (see Figure 1 for detailed protocol). Injections were administered between 9:00 a.m. and 12:00 p.m.

**Figure 1.**
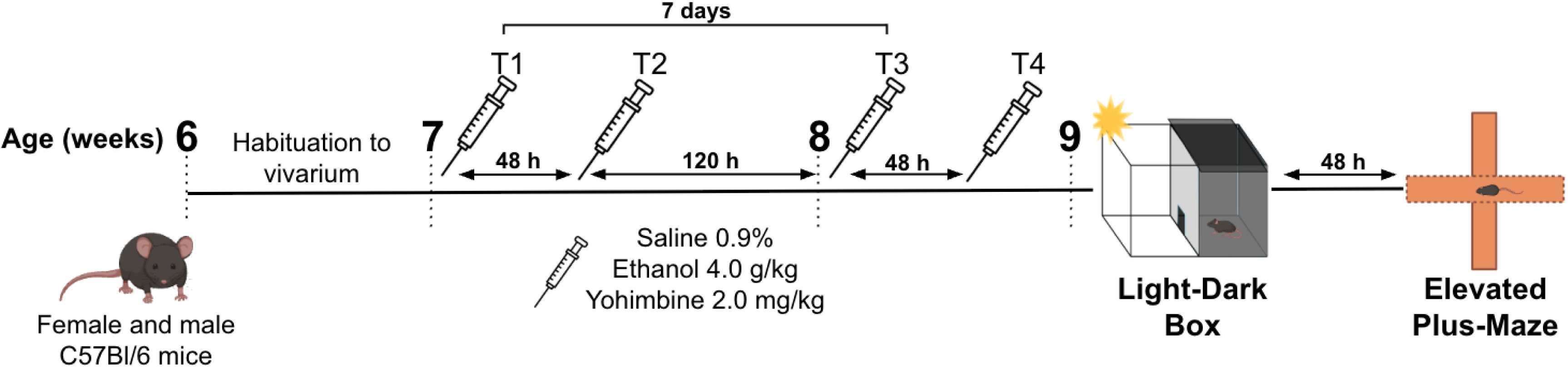
Timeline of the experimental protocol, illustrating the distribution of four administrations of ethanol, yohimbine, or saline over a two-week period. Numbers represent the age (in weeks) animals were at during each procedure.

### Blood ethanol concentration and plasma corticosterone measurements

Blood samples were collected from a separate cohort of C57BL/6 mice, distinct from those later used in the LDB and EPM experiments. Samples were obtained 30 min after i.p. injection on the first (T1) and last (T4) days of drug protocol via submandibular vein puncture using a lancet (Golde et al., 2005). Blood was stored in tubes containing ethylenediaminetetraacetic acid (EDTA; 60 mg/ml), centrifuged at 2,300 rpm at 4°C for 15 minutes, and the supernatant (plasma) was collected and stored at −20°C.

Blood ethanol concentration (BEC) analysis was performed at the Laboratory of Toxicological Information and Assistance Center (CIATox) of the Universidade de Campinas (Unicamp), using gas chromatography with a flame ionization detector (GC-FID 2010 Plus, Shimadzu, Japan) coupled to a headspace autosampler (HS-10, Shimadzu, Tokyo, Japan). The chromatographic separation was performed on a DB-BAC1 UI chromatographic column (30 m x 0.320 mm x 1.8 µm; Agilent Technologies, USA), maintained at 40°C (isothermal), with a column equilibration time of 1 min. The injection block parameters were: injector temperature at 150°C, injection mode in split mode 1:40, carrier gas pressure at 60.5 kPa, total flow of 85 mL/min, column flow of 2 mL/min, linear velocity at 33.7 cm/sec, purge flow of 3 mL/min, and flow control mode in linear velocity. The flame ionization detector parameters were: detector temperature at 250°C, flame composed of hydrogen (40 mL/min), synthetic air (400 mL/min) and makeup gas (nitrogen, 30 mL/min). The total run time was 5 min. The autosampler parameters were: oven temperature of 80°C, transfer line temperature of 105°C, sample line temperature of 90°C, pressurize gas pressure of 100 kPa, shaking level of 2, shaking time of 1 min, shaking equilibre time of 0.5 min, equilibrating time of 4 min, pressurizing time of 1 min, pressurization equilibre. time of 0.17 min, load time of 0.5 min, load equilibre time of 0.17 min, injection time of 1 min and GC cycle time of 5.6 min. The test detection range was 0.05 to 5.0 g/L.

Plasma corticosterone was analysed in duplicates using an enzyme-linked immunosorbent assay (ELISA) with a commercial kit (RE52211, IBL International, Hamburg, HH, Germany). The assay sensitivity was 0.589 nmol/L, with intra- and inter-assay mean variations of 5.15% and 7.08%, respectively.

### Behavioral procedures

We employed two well-established paradigms to assess anxiety-like behavior in rodents: the LDB and the EPM. The LDB consisted of an apparatus with two compartments: one highly illuminated (∼400 lux) (Takao and Miyakawa, 2006), and other mostly dark (∼3 lux) side. Compartments measured the same size (22 x 24 x 26 cm), and were connected by a guillotine door. The animal was placed in the dark compartment with the guillotine door closed, which was opened after 5 seconds to start the test. It is important to point out that the LDB protocol used in this work was slightly different from classic protocols, which involve a smaller dark side and initiating the test with the animal in the light side (Crawley and Goodwin, 1980; Bourin and Hascoët, 2003). Our settings facilitate the observation of risk assessment behaviors because the animal must evaluate the aversive side before exploring it.

Our EPM consisted of a wooden apparatus, elevated 50 cm above the ground, with two open and two closed (protected) arms. The open and closed arms equally measured 27.8 cm and the height of closed arms walls was 14 cm. Arms were connected by a square central area (7.8 x 7.8 cm). Luminosity reached ∼46 and ∼16 lux in the open and closed arms, respectively, similar to previous reports (Yu et al., 2007; Bruchas et al., 2009; Tseitlin et al., 2023; Pichinin et al., 2025, preprint). The maze was fully covered in waterproof varnish to prevent urine impregnation. The test started with the animal placed on the center of the maze, facing one of the open arms.

Behavioral testing initiated five to seven days after the last drug exposure. Animals were acclimated to the experiment room for 30 minutes before testing. The LDB and EPM tests were performed during the light phase of the light/dark cycle (between 12:00 and 3:00 p.m.) Tests were recorded using a webcam (Logitech C920, 30 fps) and streamed to a computer in an adjacent room. Each mouse was tested only once on each apparatus, and the apparatuses were cleaned with a 20% ethanol solution between sessions. After each session, animals were immediately placed back in their homecage. Animals were tested in the LDB and, 48 h later, in the EPM. In both tests, animals were allowed to explore the apparatus for 5 minutes.

During this time, the duration and frequency of behaviors, including time spent, number of risk assessment behaviors, entries in each compartment, as well as other behaviors listed in Table S1, were recorded. Risk assessment behaviors were evaluated into two categories: NoGo behaviors, in which the animal, right after evaluating the risk, does not enter the aversive compartment, and Go behaviors, in which the animal decides to explore the aversive environment after risk assessment (Pichinin et al., 2025, preprint). It is important to note that, due to limitations in our setup, LDB behaviors were recorded only on the light side.

### Data exportation and statistical analyses

Recorded videos of the LDB and EPM were analyzed using BORIS (v. 8.21.8), an open-source software for behavior analysis (Friard and Gamba, 2016). All behaviors listed in Table S1 were quantified by the experimenter, who was blinded to the experimental groups. Data were exported and processed using a modified version of a custom-made Python (v. 3.7.6) code (available at [GitHub](https://github.com/mariannacecyn/App-for-Behavior-Data-Processing.git)). Locomotion (distance and velocity) during the EPM test was assessed using ezTrack, an open source video analysis pipeline (Pennington et al., 2019). Go and NoGo risk assessment indexes were calculated by summing the number of Go and NoGo head outs and stretches, respectively, for each animal.

The frequency of discrete behaviors and transition probabilities were compared using Generalized Linear Models (GzLM), with Poisson and linear distributions applied as appropriate. Linear Mixed Models (LMM) were used to compare BEC levels and corticosterone levels between the first and last treatment days, with sex and treatment day as factors.

Behavioral sequences were analyzed using a first-order Markov chain framework to quantify the global contribution of specific behavioral transitions to the animals’ overall behavioral repertoire, as in Pichinin et al. (2025; preprint). For these analyses, data from female and male mice were pooled, as sex did not significantly affect the majority of behavioral variables analyzed. For each animal, the probability of occupying each behavioral state (p(A)) was computed based on the relative frequency of that behavior across the session. To capture both state occupancy and sequential organization across the entire behavioral chain, for each pair of behaviors A→B, transition values were computed by multiplying the probability of occupying the initial state by the probability of observing the subsequent behavior, yielding a joint probability metric of p(A) × p(A→B). Given the limited availability of standardized statistical approaches for direct multi-group comparisons of behavioral Markov chains, group-level analyses were performed using pairwise comparisons between the controls and each experimental group (Ctrl vs EtOH; Ctrl vs Yoh), as defined *a priori* to directly address the study’s primary hypotheses regarding the effects of alcohol exposure and pharmacological stress on behavioral organization. In line with previous studies (Espejo, 1997; Lino-de-Oliveira et al., 2005), state events occurring less than 1% of the time and transitions with less than 2% occurrence were excluded from the analyses.

To assess relationships between variables, we ran correlation analyses. A Shapiro-Wilk normality test was first performed and, because not all variables followed a normal distribution, Spearman’s rank correlation was used. Analyses were conducted with animals separated by experimental groups to determine whether relationships between variables were group-dependent. Correlations were performed without considering sex, as this factor did not have a significant effect on the majority of variables analyzed using GzLM.

Markov chains were built using Microsoft PowerPoint (v. 2411), with circles proportional to the frequency of each behavior, and arrow transparency proportional to transition probabilities. Bonferroni post hoc tests were applied when necessary, and a p-value < 0.05 was considered statistically significant. Statistical analyses were performed using Jamovi (v. 2.4.8.0) and graphs were assembled in Graphpad Prism (v. 10.1.1). Data are presented as mean ± standard error of the mean (SEM), with all statistical details provided in the figure legends.

## Results

### Ethanol administration induces high levels of intoxication and sustained elevated levels of corticosterone, similar to yohimbine

Ethanol (4.0 g/kg) induced high levels of BEC that remained stable across treatment days and did not differ between females and males (Figure 2A). Ctrl mice exhibited a significant decrease in corticosterone response over the protocol period, with lower levels on T4 compared to T1 (p = 0.014), suggesting HPA axis adaptation to repeated handling and injections (Figure 2B). In contrast, EtOH and Yoh effects on corticosterone persisted over the two-week treatment period, indicating sustained activation of the HPA axis relative to controls.

**Figure 2.**
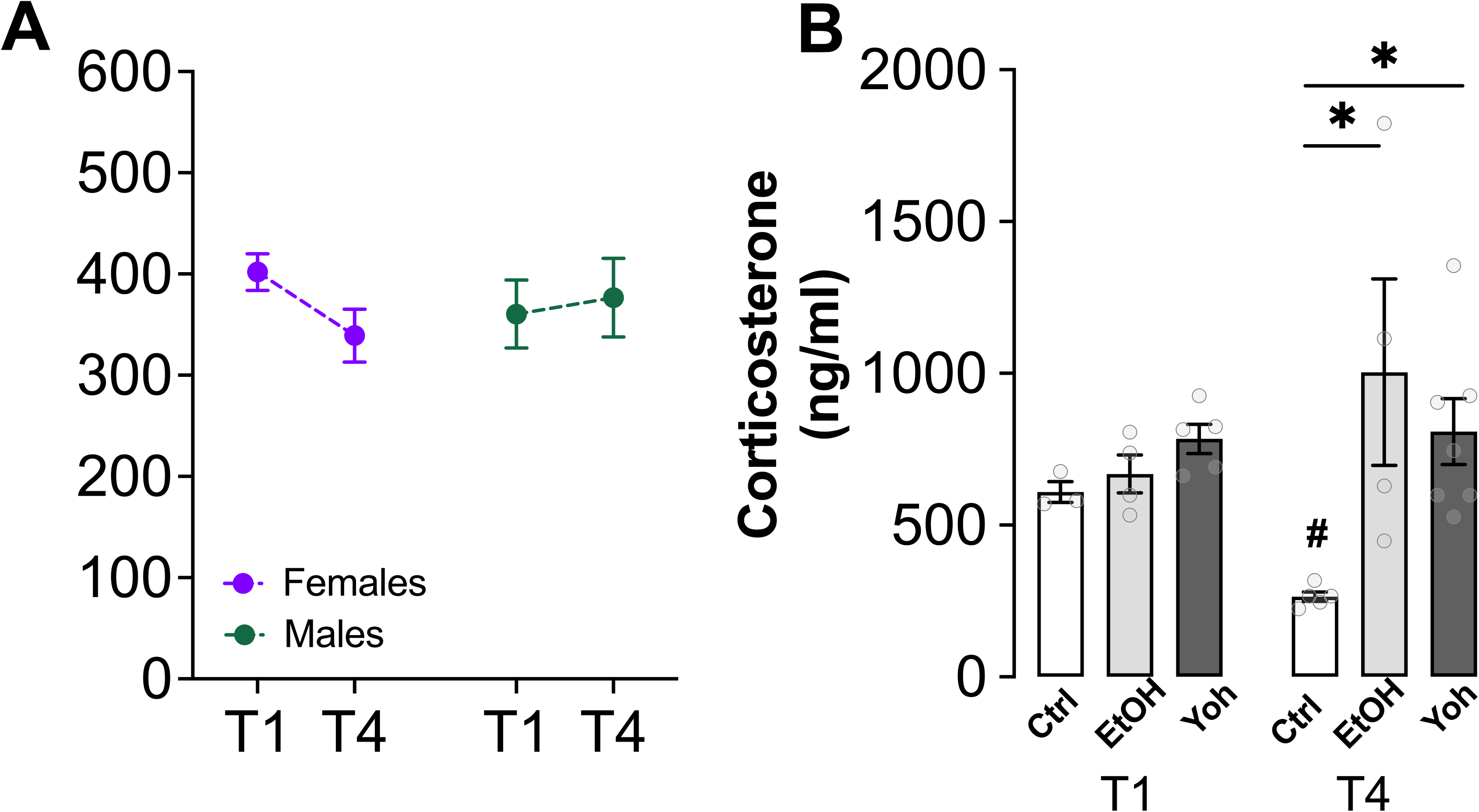
Blood ethanol concentrations (BEC) and plasma corticosterone measurements following ethanol (EtOH) and yohimbine (Yoh) administrations. **(A)** Blood EtOH Concentrations (BEC) 30 minutes after i.p. administration on the first (T1) and last (T4) days of the protocol. Mice presented similar levels of BEC on the first and last treatment days (F_(1, 4)_ = 0.69). Sex was not significant (F_(1, 4)_ = 0.03). Sex-separated analyses showed similar levels of BEC on the first and last treatment days in females (F_(1, 2)_ = 1.60) and males (F_(1, 2)_ = 3.00). n = 6/sex. **(B)** Plasma corticosterone levels on the first and last days of treatment. Groups presented similar levels of plasma corticosterone on T1, whereas on T4 corticosterone levels were significantly higher in the EtOH and Yoh groups in comparison to the control (Ctrl) group (treatment day: F_(1, 16)_ = 0.01; group: F_(2, 16)_ = 8.77, p = 0.003, group*treatment day: F_(2, 16)_ = 4.70, p = 0.025. * p < 0.05 in comparison to Ctrl on T4; # p < 0.05 in comparison to the same group on T1). Ctrl: n = 4/group; EtOH: N= 4/group; Yoh: n = 6/group.

### Ethanol and yohimbine exposures during late adolescence impair risk assessment behaviors in adulthood in both female and male mice

To test whether ethanol intoxications or stress exposures during late adolescence (7-8 weeks of age) could affect behavior in early adulthood (9 weeks of age), mice were tested in the LDB five days after the last drug exposure. Latency to pass to the light side was similar in all groups, regardless of sex (Figure 3A). The percentage of time spent in the light side was similar among the groups in female mice, but ethanol-treated male mice spent less time in the light in comparison to controls and yohimbine-treated mice (EtOH vs. Ctrl: p = 0.03; EtOH vs. Yoh: p = 0.035) (Figure 3B). These results indicate that late adolescence ethanol intoxication episodes, but not yohimbine exposures, induce a significant anxiogenic behavior in males but not in females. No effects on transitions between the dark and light sides were observed (Figure 3C), although intoxication with ethanol influenced other exploratory behaviors in the LDB. For instance, ethanol-treated males performed less rears compared to the other groups (EtOH vs Ctrl: p = 0.009, EtOH vs. Yoh: p = 0.011). Sniffs and other exploratory behaviors were not affected by ethanol or yohimbine exposures (Figure S1A-C). A central goal of this study was to determine whether prior alcohol intoxication or stress exposure can promote risky decision-making later in life. To address this, we quantified risk assessment behaviors, distinguishing between NoGo responses, when animals assess risk but choose not to enter the aversive environment, and Go responses, in which mice decide to move toward the aversive compartment after evaluating risk. Ethanol and stress exposures induced fewer NoGo risk assessment when compared to control mice in both sexes (females: EtOH vs Ctrl: p = 0.015, Yoh vs. Ctrl: p = 0.027; males: EtOH vs Ctrl: p < 0.001, Yoh vs. Ctrl: p < 0.001). On the other hand, Go risk assessment was not affected by prior treatment in both sexes (Figure 3E). To better characterize how often animals chose the risky option (Go decisions), we compared the proportion of Go risk assessment events relative to the total number of risk-assessment behaviors (Figure 3F). There was no significant effect of sex. Control female mice predominantly returned to the dark side after assessing risk, whereas ethanol-treated mice more frequently transitioned to the light side (EtOH vs. Ctrl: p = 0.028). A similar pattern was observed in males, in which both ethanol- and yohimbine-treated animals showed a higher proportion of Go events compared with controls (EtOH and Yoh vs. Ctrl: p < 0.001).

**Figure 3.**
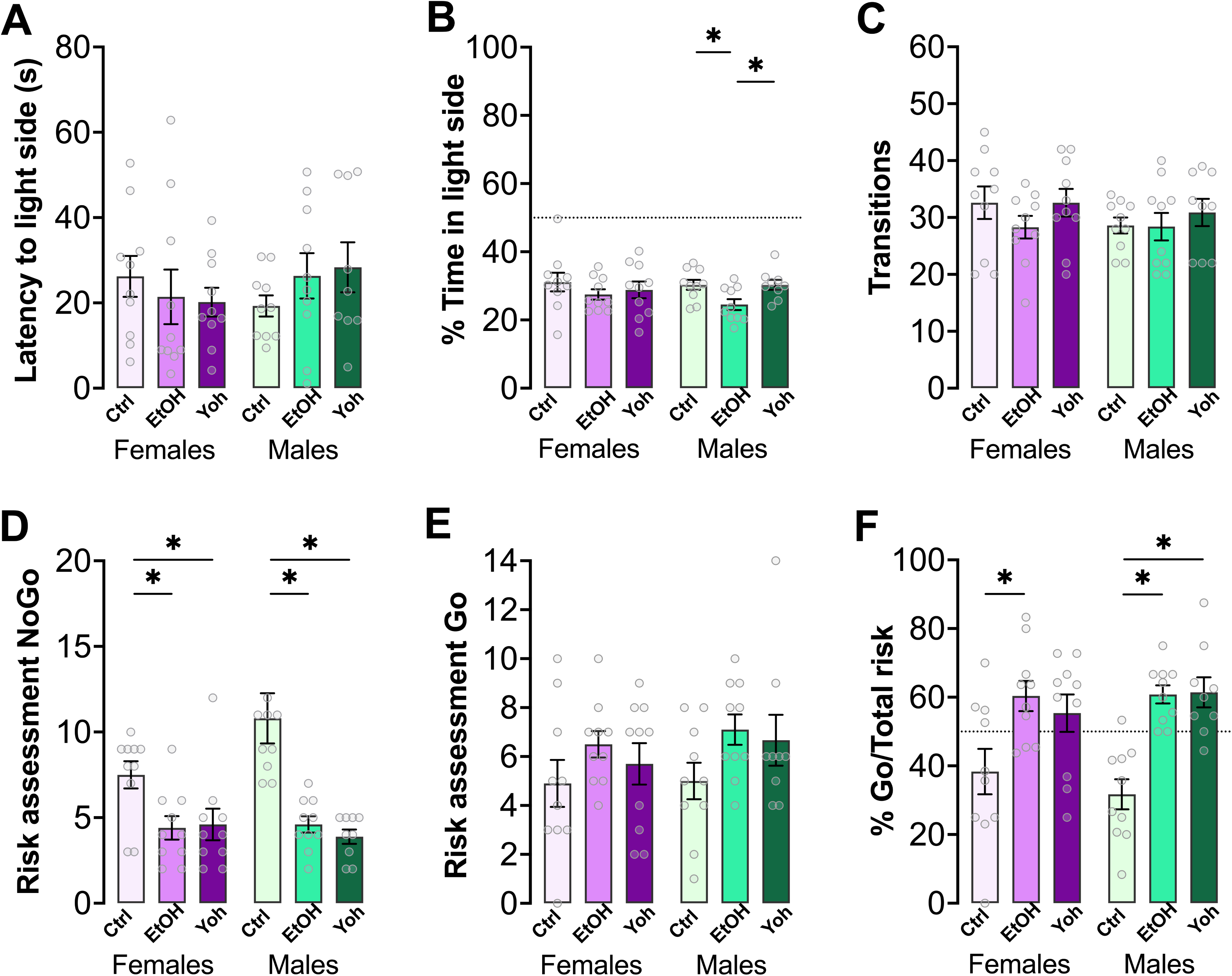
Anxiety-like, exploratory and risk assessment behaviors of adult female and male mice (9 weeks old) in the LDB five days after saline (Ctrl), ethanol (EtOH) or yohimbine (Yoh) exposures during late adolescence. **(A)** Latency to pass to the light side was similar in both sexes (X²_(1)_ = 0.37). Sex-separated analyses showed no group effect (females: X²_(2)_ = 0.80; males: X²_(2)_ = 2.06). **(B)** Percentage of time in the light side was similar in both sexes (X²_(1)_ = 0.24). Sex-separated analyses showed that males in the EtOH group spent less time in the light side compared to Ctrl and Yoh (X²_(2)_ = 10.10, p = 0.006; EtOH vs. Ctrl: p = 0.03, EtOH vs. Yoh: p = 0.035). No significant group differences were found in females (X²_(2)_ = 1.28). **(C)** Transitions were similar in both sexes (X²_(1)_ = 1.74). Sex-separated analysis showed no effect of group (females: X²_(2)_ = 4.02; males: X²_(2)_ = 1.21). **(D)** NoGo risk assessment analysis showed no significant effect of sex (X²_(1)_ = 0.58). Sex-separated analysis showed a significant effect of group for females (X²_(2)_ = 10.40, p = 0.005; EtOH vs. Ctrl: p = 0.015, Yoh vs. Ctrl: p = 0.027) and males (X²_(2)_ = 40.90, p < 0.001; EtOH and Yoh vs. Ctrl: p < 0.001). **(E)** Go risk assessments were similar in both sexes (X²_(1)_ = 0.71). Sex-separated analyses showed no significant group effect (females: X²_(2)_ = 2.25; males: X²_(2)_ = 4.04). **(F)** The percentage of Go risk assessment among total risk assessment behaviors was similar in both sexes (X²_(1)_ = 0.00). Sex-separated analysis showed significant group effects for females (group: X²_(2)_ = 8.63, p = 0.013; EtOH vs. Ctrl: p = 0.028) and males (group: X²_(2)_ = 39.40, p < 0.001; EtOH and Yoh vs. Ctrl: p < 0.001). The dashed line represents 50%, indicating an equal distribution between Go and NoGo risk assessment. Ctrl: n = 10/sex; EtOH: n = 10/sex; Yoh: n = 10 females, 9 males.

To analyse first-order transitions within the ethogram, we calculated the transition probabilities to and from each behavior and constructed Markov chains for each group illustrating the resulting behavioral structure during the LDB test (Figure 4). For this analysis, data from male and female mice were pooled, as frequentist analyses of discrete behaviors revealed no robust sex-dependent differences, and pooling increased both statistical power and the stability of transition probability estimates. Briefly, animals exposed to ethanol and yohimbine showed significantly fewer transitions from the dark compartment to NoGo risk assessment behaviors compared to controls (i.e., dark entry - NoGo head out, EtOH vs. Ctrl: p = 0.002; Yoh vs. Ctrl: p = 0.005). In contrast, these animals exhibited increased transitions toward Go risk assessment behaviors (i.e., dark entry - Go stretch, EtOH vs. Ctrl: p = 0.004; Yoh vs. Ctrl: p = 0.010 and dark entry - Go head out, EtOH vs. Ctrl: p = 0.002), indicating a shift toward less cautious exploratory behavior. Please refer to Table S2 for a complete list and statistical description of significant differences in transition probabilities between the EtOH and Yoh groups compared with Ctrl.

**Figure 4.**
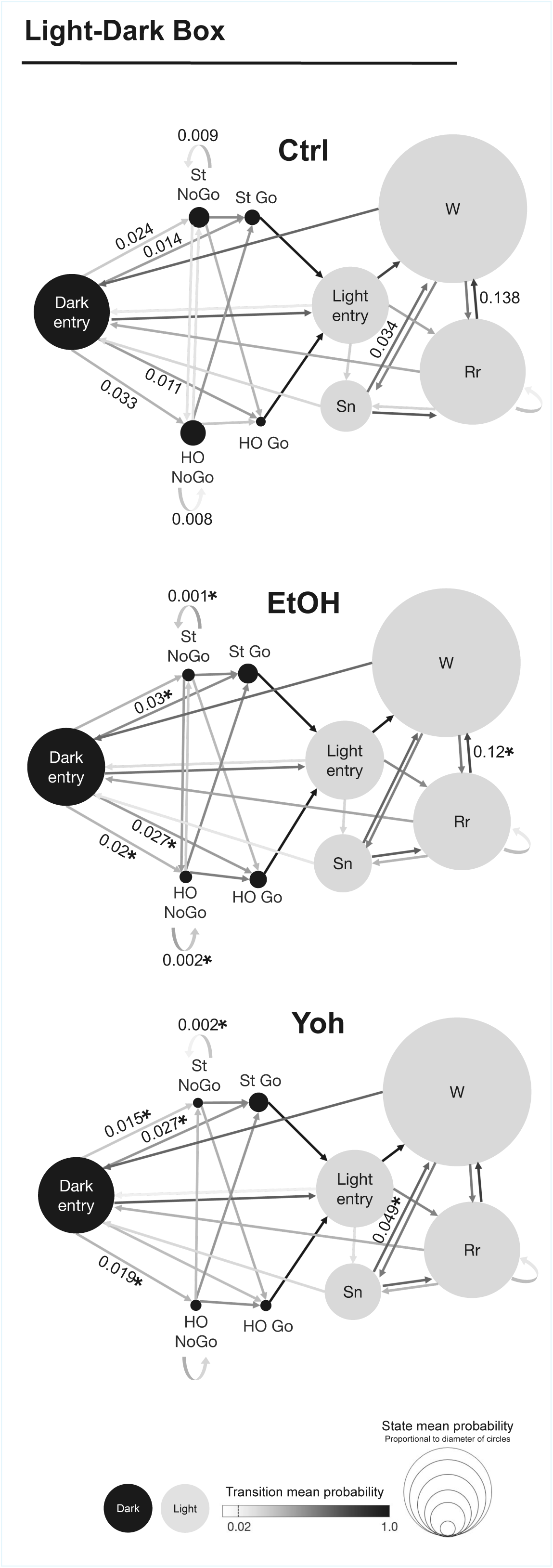
Markov chains representing the sequencing and chaining of behavioral events in the experimental groups during the LDB test. Black circles represent behaviors in the dark side and gray circles represent behaviors in the light side. Circles’ sizes are proportional to the number of occurrences of each behavior, while the arrows’ color gradient represents the probability of each transition. Numbers represent transition probabilities. For experimental groups, only transitions significantly different from Ctrl are displayed, with asterisks indicating p < 0.05. Complete statistics for these transitions are listed in Table S2. HO: head out, St: stretch, Rr: rear, Sn: sniff, W: walk. Ctrl: n = 20 (10/sex); EtOH: n = 20 (10/sex); Yoh: n = 19 (10 females, 9 males).

Seven days after the last drug exposure, and two days after LDB, animals were tested on the EPM. Anxiety-like and exploratory behaviors in the EPM were not affected by ethanol and yohimbine exposures during late adolescence, as observed in sex-separated analyses (Figure 5A-B; Figure S2A-F). The number of transitions was similar in all groups, regardless of sex (Figure 5C), as well as distance and velocity throughout EPM testing (Figure S3A-B). In contrast, both ethanol and yohimbine reduced the frequency of NoGo risk assessment behaviors similarly in females (EtOH vs. Ctrl: p = 0.009, Yoh vs. Ctrl: p = 0.002) and males (EtOH vs. Ctrl: p < 0.001; Yoh vs. Ctrl: p = 0.006; Figure 5D). Go risk assessment behaviors were similar in all groups, regardless of sex (Figure 5E). Additionally, there was no effect of sex on the proportion of Go risk assessment behaviors among total risk assessment behaviors, and group effects were not not observed in sex-separated analyses (Figure 5F).

**Figure 5.**
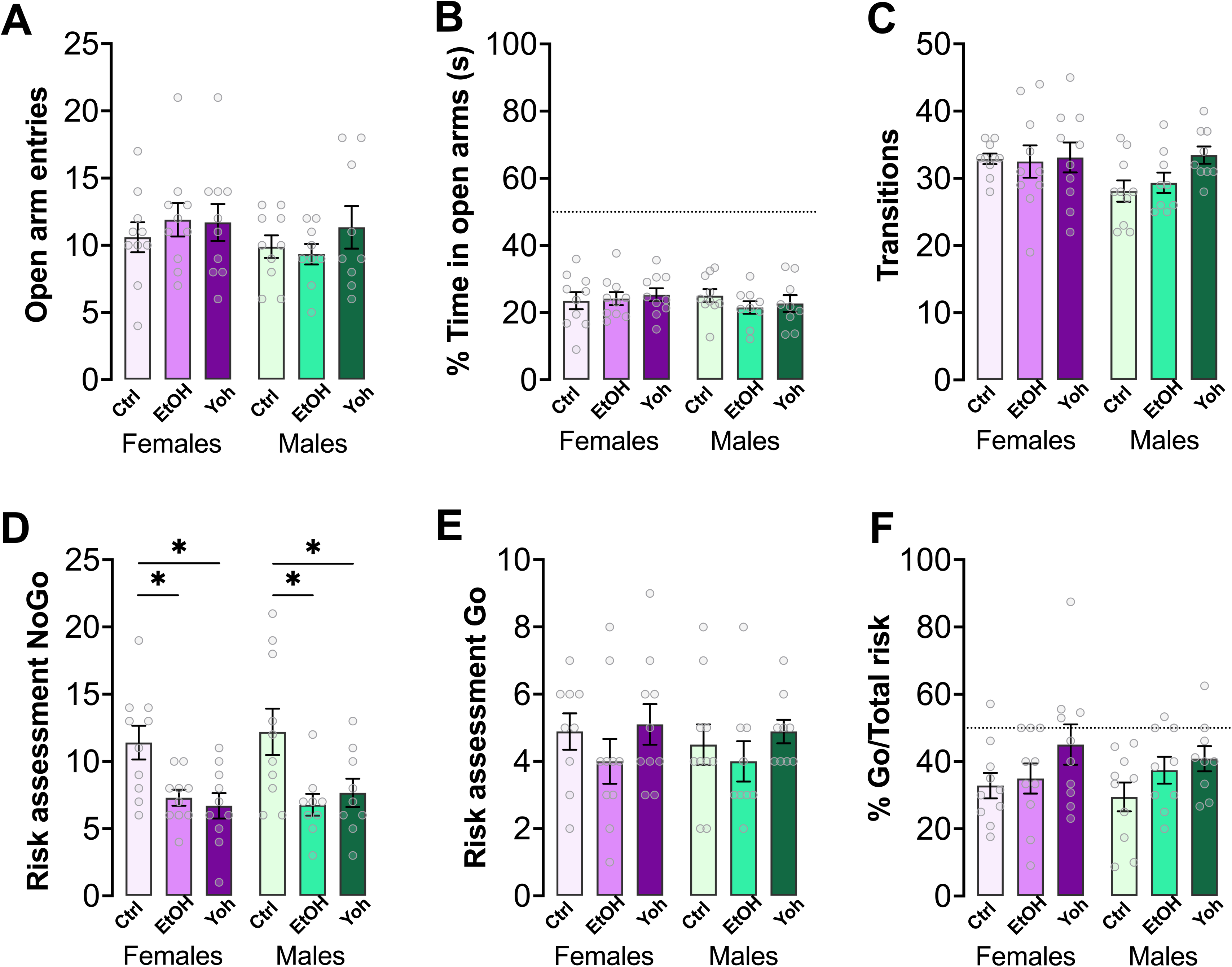
Anxiety-like, exploratory and risk assessment behaviors of adult female and male mice (9 weeks old) in the EPM seven days after saline (Ctrl), ethanol (EtOH) or yohimbine (Yoh) exposures during late adolescence. **(A)** Number of open arm entries was similar in both sexes (X²_(1)_ = 1.75). Sex-separated analyses showed no group effects (females: X²_(1)_ = 0.87; males: X²_(1)_ = 1.86). **(B)** Percentage of time in the open arms was similar in both sexes (X²_(1)_ = 0.42). Sex-separated analyses showed no group effects (females: X²_(1)_ = 0.34; males: X²_(1)_ = 1.50). **(C)** Transitions were similar in both sexes (X²_(1)_ = 2.71). Sex-separated analyses showed no group effects (females: X²_(1)_ = 0.06; males: X²_(1)_ = 4.75). **(D)** NoGo risk assessment was similar in both sexes (X²_(1)_ = 0.13). Sex-separated analysis showed similar group effects in females (X²_(2)_ = 14.80, p < 0.001; EtOH vs. Ctrl: p = 0.009, Yoh vs. Ctrl: p = 0.002) and males (X²_(2)_ = 17.50, p < 0.001; EtOH vs. Ctrl: p < 0.001; Yoh vs. Ctrl: p = 0.006). **(E)** Go risk assessment was similar in both sexes (X²_(1)_ = 0.38). Sex-separated analyses showed no group effects (females: X²_(2)_ = 2.80; males: X²_(2)_ = 1.76). **(F)** The percentage of Go risk assessment among total risk assessment behaviors was similar in both sexes (X²_(1)_ = 0.12). Sex-separated analysis did not point to significant group effects (females: X²_(2)_ = 3.62; males: X²_(2)_ = 4.26). The dashed line represents 50%, indicating an equal distribution between Go and NoGo risk assessment. Ctrl: n = 10/sex; EtOH: n = 10 females, 9 males; Yoh: n = 10 females, 9 males.

To examine the structural organization of behaviors in the EPM, we analyzed transition probabilities between exploratory and risk assessment behaviors and constructed Markov chains summarizing the behavioral structure of the task (Figure 6). In this context, ethanol- and yohimbine-exposed animals exhibited fewer transitions from walking in the closed arms to NoGo head outs compared to controls (EtOH vs. Ctrl: p = 0.016; Yoh vs. Ctrl: p = 0.001), indicating a reduced engagement in cautious risk assessment from protected areas. Additionally, experimental groups showed lower transition probabilities from sniffing to rearing relative to controls (EtOH vs. Ctrl: p = 0.017; Yoh vs. Ctrl: p = 0.04), which is consistent with diminished vertical exploration. Other significant differences from EtOH and Yoh groups to Ctrl are enlisted in Table S3. Together, these alterations suggest that ethanol and yohimbine modify the structure of behavior in the EPM, altering how exploratory and risk assessment events are sequentially organized.

**Figure 6.**
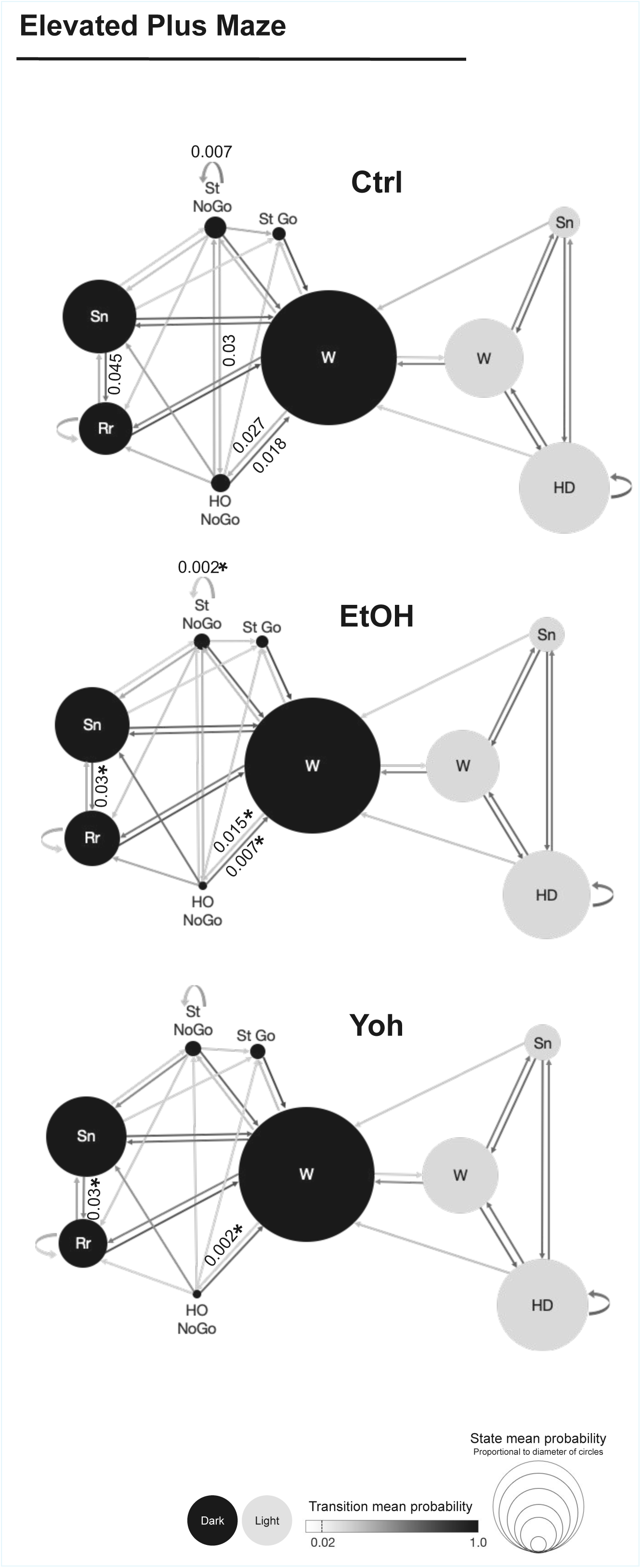
Markov chains representing the sequencing and chaining of behavioral events in the experimental groups during the EPM test. Black circles represent behaviors in the closed arms and gray circles represent behaviors in the open arms of the maze. Circles’ sizes are proportional to the number of occurrences of each behavior, while the arrows’ color gradient represents the probability of each transition. Numbers represent transition probabilities. For experimental groups, only transitions significantly different from Ctrl are displayed, with asterisks indicating p < 0.05. Complete statistics for these transitions are listed in Table S3. HO: head out, St: stretch, Rr: rear, Sn: sniff, W: walk. Ctrl: n = 20 (10/sex); EtOH: n = 19 (10 females, 9 males); Yoh: n = 19 (10 females, 9 males).

To further characterize how individual behavioral measures relate to one another beyond transition structure, we next examined pairwise correlations among behavioral variables. Data from female and male mice were also pooled for correlation analyses. In the LDB test, the control group showed a structured pattern in which a classical anxiety-related measure (represented by the percentage of time in the light compartment) was positively correlated with exploratory behaviors such as transitions and rears, while risk assessment behaviors were weakly correlated or largely independent from exploratory measures (Figure 7A). Following ethanol exposure, this structure was altered, with stronger positive correlations emerging between exploratory behaviors, alongside with negative correlations between NoGo risk assessment (in particular, NoGo stretches) and anxiety-like and exploratory behaviors. Interestingly, ethanol-exposed animals showed a negative correlation between the two categories of Go risk assessment behaviors (Figure 7B). These patterns indicate a reorganization of behavioral relationships toward increased engagement with the aversive compartment. In contrast, yohimbine exposure produced a distinct correlation pattern characterized by strong negative correlations between exploratory measures and NoGo head outs, indicating that ethanol and yohimbine can differentially affect risk assessment behavioral nuances (Figure 7C).

**Figure 7.**
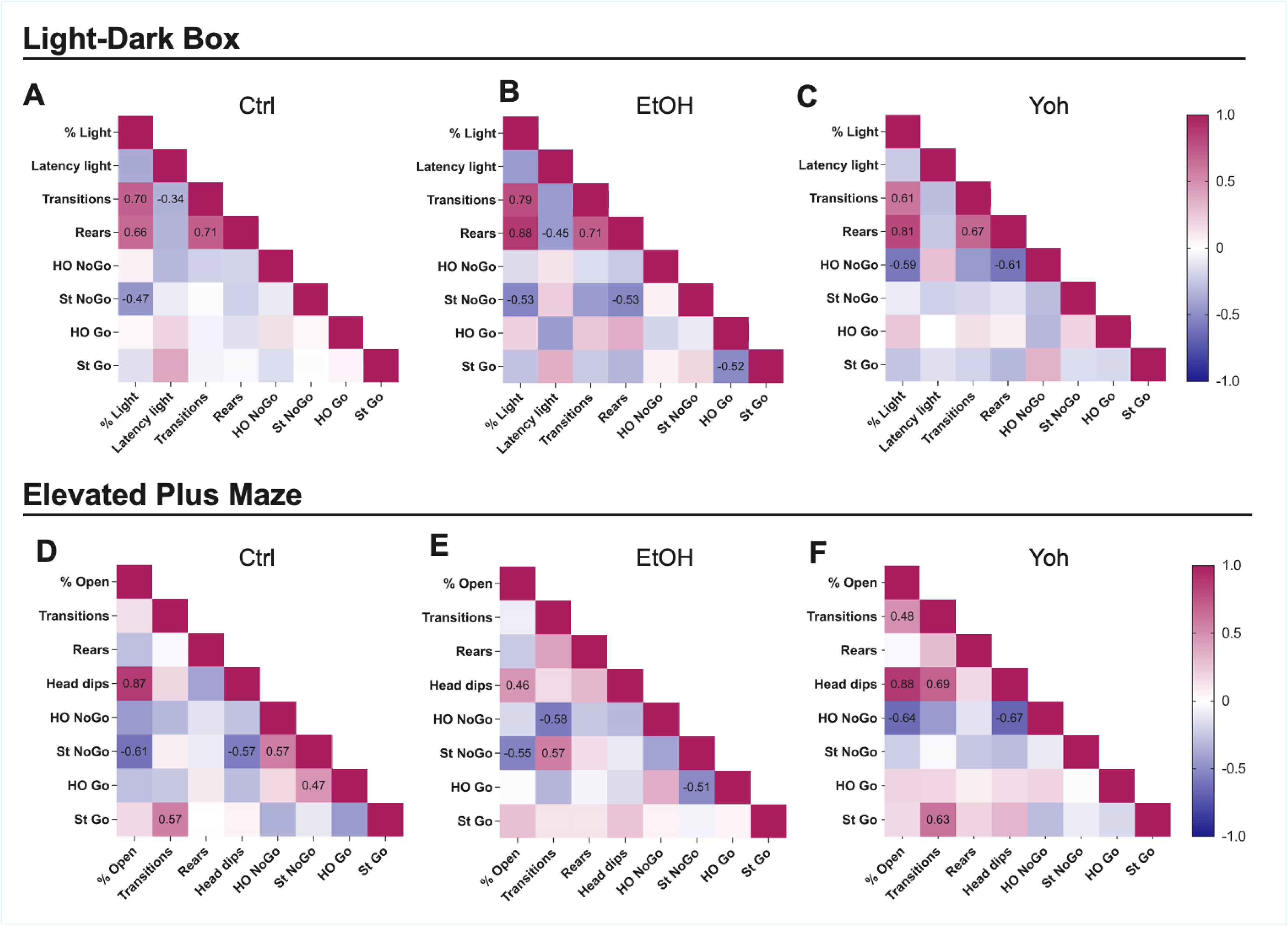
Spearman’s correlation heat maps for female and male pooled data from the LDB and EPM tests, respectively. Heatmaps depict Spearman’s rank correlation coefficients (ρ) between exploratory and risk assessment behaviors for control (Ctrl), ethanol- (EtOH), and yohimbine-exposed (Yoh) mice. Numerical values shown within the cells correspond to statistically significant correlations only (Spearman’s ρ, p < 0.05). Color intensity reflects the strength and direction of the correlation. Non-significant correlations are left blank. Data from male and female mice were pooled for correlation analyses.

EPM correlation analyses revealed that control animals exhibited strong positive correlations between exploratory behaviors, such as head dips, and anxiety-like behavior, represented by the percentage of time in the open arms. Notably, NoGo risk assessment was negatively correlated to head dips (Figure 7D). In ethanol-exposed mice, this pattern was altered, with a weakened association between exploratory and anxiety-like behaviors (Figure 7E). In contrast, yohimbine-treated animals displayed stronger positive correlations between exploratory and anxiety-like measures, including transitions, head dips, and percentage of time in the open arms, as well as and stronger negative correlations between NoGo risk assessment (particularly NoGo head outs), and both anxiety-like and exploratory behaviors (Figure 7F).

## Discussion

The present study demonstrates that both repeated ethanol intoxication or pharmacological stress repeated exposure during late adolescence produce persistent alterations in risk assessment behavior in early adulthood. Both treatments led to a marked reduction in NoGo risk assessment behaviors in the LDB and the EPM, as well as changes in behavior structure, shown by differences in the transition probabilities among behavioral events and the correlations among discrete behaviors. These results indicate a bias toward less cautious decision-making when animals previously exposed to ethanol and yohimbine are presented with new, potentially threatening environments.

Risk assessment, a defensive strategy animals use to balance exploration of unfamiliar environments with the need to avoid potential threats, is reliably evoked in approach-avoidance paradigms such as the LDB and the EPM (Augustsson and Meyerson, 2004). Importantly, risk assessment behaviors can be modulated independently of general anxiety levels, providing information that is not captured by traditional measures such as time spent in open arms or illuminated compartments. Moreover, to investigate risk assessment behaviors more thoroughly, they can be classified according to the animal’s level of exposure to the aversive context (i.e., exposing only the head or half its body) or according to the decision made after assessing the risk (i.e., exploring the open arms/light side or not; Pichinin et al., 2025, preprint).

In the present study, exposure to high doses of ethanol or the α2-adrenergic antagonist yohimbine during late adolescence biased animals toward risk-taking behaviors in adulthood, as reflected in decreases of NoGo risk assessment, with no major effects on general exploration or anxiety-like behaviors. These results suggest a diminished tendency to withhold action following risk assessment, reflecting a shift toward riskier decision strategies. There is substantial evidence that both alcohol and stress influence impulsivity, risk assessment, and other aspects of decision-making. In humans, acute alcohol consumption reduces inhibitory control (Gan et al., 2014; Weafer et al., 2021) and increases impulsivity (Reynolds et al., 2006; Dougherty et al., 2008).

Weafer et al. (2021) found that neural correlates of inhibitory control were associated with stimulant-like responses to alcohol, indicating that alcohol’s effects on brain activity can underlie impairments in the ability to suppress actions. A meta-analysis by McPhee and Hendershot (2023) further confirmed that acute alcohol consistently impairs the ability to withhold or stop actions in Go/NoGo and stop signal tasks, with stronger effects at higher intoxication levels and in conditions requiring a strong prepotent response, highlighting its robust impact on inhibitory control and its contribution to impulsive behavior. Parallel findings in rodents indicate that ethanol produces lasting behavioral and prefrontal alterations that persist beyond acute intoxication. In rats, intraperitoneal ethanol administrations during adulthood induced impairments in reversal learning that were evident during abstinence (5-7 days after last exposure), at the same dose used in the present study (Badanich et al., 2016). Consistently, chronic alcohol administered in adult mice leads to enduring alterations in medial prefrontal cortex plasticity, including increased NMDA/AMPA ratios, structural spine remodeling, and attention deficits that persist for at least one week into withdrawal, long-lasting disruptions of prefrontal-dependent executive functions (Kroener et al., 2012). These findings support the notion that repeated alcohol impairs inhibitory control and risk assessment, providing a framework to interpret the reductions in NoGo behaviors observed in the present study.

Stress can also disrupt inhibitory control and risk assessment. Acute stress impairs inhibitory control and decision-making in humans, with individual susceptibility influenced by autonomic regulation and anxiety levels (Roos et al., 2017; Van den Bussche et al., 2020). In rats, stress selectively alters cost/benefit decision-making, influencing the balance between cautious and risky choices (Shafiei et al., 2012). Pharmacological stressors such as yohimbine increase risky choice in risk discounting tasks when administered shortly before testing, an effect linked to heightened catecholaminergic signaling within prefrontal circuits (Münster et al., 2024). By blocking α2-adrenergic autoreceptors, yohimbine enhances noradrenergic tone in limbic and prefrontal regions involved in arousal, emotional processing, and behavioral inhibition, promoting anxiogenic-like states and reduced cautious evaluation (Blanchard et al., 1993; Tanaka et al., 2000; Vasa et al., 2009; Sperl et al., 2022). These findings provide a basis to interpret the decreases in NoGo behaviors following repeated yohimbine exposures in the present study. Additionally, our results align with previous findings showing an increased proportion of open arm entries following stretch-attend postures in the closed arms of the EPM in rats exposed to chronic social stress (Stickling and Rosenkranz, 2025). Together, these findings suggest that repeated activation of stress systems during adolescence, whether driven by intoxication or pharmacological challenge, can produce lasting biases in risk assessment strategies. Notably, the present study extends this literature by showing that repeated episodes of heavy ethanol intoxication and yohimbine during late adolescence bias adult behavior toward reduced defensive evaluation, even when testing occurs days after the last exposure.

Age, sex and dosage are critical biological and procedural variables that shape the long-lasting effects of alcohol or stress exposures. Adolescence represents a critical developmental window for the maturation of neural circuits supporting behavioral inhibition, risk assessment, and decision-making. Alcohol exposure during this period has been shown to produce lasting alterations in risk-related strategies and inhibitory control, effects that persist into adulthood (Towner and Varlinskaya, 2020). Previous work from our group has indicated that a repeated moderate dose of ethanol (3.2 g/kg) during early adolescence (weeks 5-6) selectively decreases NoGo risk assessment in adult male mice submitted to LDB testing (Pichinin et al., 2025, preprint). Extending these findings to late adolescence (weeks 7-8), the present study shows that higher-dose ethanol exposures, as well as pharmacologically induced stress, reduce NoGo risk assessment in adult male and female mice in both the LDB and EPM. Thus, late adolescence may represent a particularly vulnerable period for disruptions in inhibitory control, resulting in a stronger tendency toward exploration and risk-taking. These results align with previous findings showing that intermittent ethanol exposure during late adolescence, at lower (2.0 g/kg) and more frequent doses than the one used in the present study, leads to lasting and timing-dependent disruptions in attention, impulsivity, and risky decision-making in mice (Sanchez-Roige et al., 2014). Our results add to this by showing that not only ethanol, but also yohimbine, reduce cautious risk evaluation, pointing to overlapping vulnerabilities in behavioral inhibition and decision-making circuits.

Beyond changes in the frequency of behaviors, our findings indicate that adolescent ethanol and stress exposures also alter how exploratory and risk assessment behaviors are sequentially organized. Using transition-based analyses and Markov chain modeling, we observed that ethanol- and yohimbine-treated animals exhibited reorganized behavioral sequences during exploration in both the LDB and the EPM. In the LDB, these alterations were characterized by reduced transitions from the protected dark compartment to NoGo risk assessment behaviors and a relative bias toward transitions leading to Go behaviors, indicating a shift in how animals engage with and act upon potentially dangerous, aversive zones. In the EPM, a complementary pattern emerged, with ethanol- and yohimbine-treated animals showing fewer transitions from walking in the closed arms to NoGo head outs, alongside altered exploratory sequences involving sniffing and rearing. Notably, these effects reflect changes in behavioral structure rather than just increases or decreases in individual behaviors, highlighting that decision-making alterations may be embedded in the temporal, sequential flow of actions.

Some studies have shown a correlation between risk assessment and anxiety-like behaviors in rodents. For example, there is evidence of increased risk assessment in animals displaying increased anxiety-like behaviors in the EPM (Ohl et al, 2001). In accordance, risk assessment behaviors are decreased by anxiolytic drugs on the same test, and also in the LDB (Cruz et al., 1994; Griebel et al., 1998). On the other hand, risk assessment behaviors can be present regardless of the level of anxiety, reflecting partially dissociable defensive strategies (Hefner and Holmes, 2007; Blanchard and Meyza, 2019). In our results, adolescent ethanol and stress exposure altered the structural integration of behavioral components rather than simply their frequency. Specifically, in both behavior tests, ethanol or yohimbine exposure led to a marked reorganization of the correlational structure of behavior linking exploratory, anxiety-like, and risk assessment behaviors relative to controls, revealing test-specific alterations in behavioral integration. In the LDB, ethanol exposure strengthened positive correlations among exploratory behaviors while promoting negative associations between NoGo risk assessment and both exploratory and anxiety-like measures. Additionally, the two categories of Go risk assessment became negatively correlated, suggesting mutually exclusive engagement of Go strategies and increased interaction with the aversive compartment. Yohimbine-treated animals showed a distinct profile characterized by strong negative correlations between exploratory behaviors and NoGo risk assessment, indicating a selective strengthening of the antagonism between exploration and cautious evaluation. A similar dissociation emerged in the EPM, where ethanol exposure weakened associations between exploratory and anxiety-like behaviors, whereas yohimbine treatment enhanced positive correlations between these measures while reinforcing negative correlations involving NoGo risk assessment. In contrast, control animals across both tests exhibited a more organized and balanced behavioral structure, with coherent covariation among exploratory and anxiety-like measures and negative associations between exploration and NoGo risk assessment, consistent with a dynamic framework in which risk assessment constrains exploration in anxiogenic environments (Rodgers and Dalvi, 1997; Ohl et al., 2001).

Some limitations to our study should be considered. Yohimbine represents a pharmacological stressor and may not fully capture the complexity of naturalistic stress experiences. Additionally, while corticosterone measurements indicate sustained HPA axis engagement, the neural mechanisms linking adolescent stress or intoxication to adult risk assessment deficits were not directly examined in this study. Future work integrating behavioral, hormonal, and circuit-level approaches will be necessary to delineate the pathways through which adolescent experiences shape adult decision-making under risk.

In summary, our findings demonstrate that repeated ethanol intoxication or stress exposure during late adolescence leads to persistent impairments in risk assessment behavior, characterized by reduced cautious evaluation and altered behavioral sequencing in adulthood. These results highlight adolescence as a critical window during which alcohol and stress can bias decision-making strategies, with potential implications for understanding the long-term behavioral consequences of early-life exposures.

## Financial support

This work was supported by the Fundação de Amparo à Pesquisa do Estado de São Paulo (FAPESP grants #2019/01686-0) and the Coordenação de Aperfeiçoamento de Pessoal de Nível Superior (CAPES, student grant #88887.695072/2022-00).

## Declaration of competing interests

The authors report no conflicts of interest.

## Acknowledgments

We thank Dr. José Luiz da Costa and Isadora Locilento Denkena (Centro de Informação e Assistência Toxicológica, Universidade Estadual de Campinas) for conducting BEC measurements. We are also grateful to Beatriz de Petribu da Costa Nunes Borges and Leticia Pichinin de Souza for their assistance with blood collection, to Andrea Aurelio Borges for her support with the plasmatic corticosterone assay, and to Marianna Nogueira Cecyn for developing the initial version of the Python code used to process behavioral data and for the assistance with the first-order Markov chain analyses. We further acknowledge Associação Fundo de Incentivo à Pesquisa (AFIP) for infrastructure support.

## Data availability

Supporting data are available from the corresponding author on reasonable request.

## Declaration of generative AI in the writing process

During the preparation of this work, the authors used ChatGPT in order to assist with English grammar only. After using this tool/service, the authors reviewed and edited the content as needed and take full responsibility for the content of the publication.

## Notes

### Competing Interest Statement

The authors have declared no competing interest.

