## Supplementary Material for "Alcohol or stress exposure during late adolescence impairs risk assessment later in life, disrupting the ethological structure of behavior in mice"

<u>**Supplemental information**</u>

[1] Programa de Pós-Graduação em Psicobiologia, Departamento de Psicobiologia, Escola Paulista de Medicina, Universidade Federal de São Paulo, São Paulo, São Paulo, Brasil.

[2] Departamento de Psicobiologia, Escola Paulista de Medicina, Universidade Federal de São Paulo, São Paulo, São Paulo, Brasil.

[3] Departamento de Fisiologia, Escola Paulista de Medicina, Universidade Federal de São Paulo, São Paulo, São Paulo, Brasil.

* Correspondent author

**Table S1:** List of analyzed behaviors during the LDB and EPM tests.

| BEHAVIOR | CATEGORY | DEFINITION |
|---|---|---|
| Rear | Exploratory | The mouse assumes a vertical position supporting itself on its hind legs while the front legs either move in the air or lean against a wall. |
| Head dip | Exploratory | Downward visual screening that happens when the mouse is in one of the elevated open arms of the plus maze. |
| Walk | Exploratory | The mouse moves around and changes position. Includes non-specific body movements such as turning or running across the apparatus. |
| Sniff | Exploratory | Rhythmic inhalation and exhalation of air through the nose while the mouse stands still. |
| Grooming* | Self-care | The mouse rubs any part of the face with circular movements of the forepaws, licks its body and/or its forepaws and back paws. |
| Light side entry | Exploratory | The mouse enters the light side of the LDB with its four paws. |
| Dark side entry | Exploratory | The mouse enters the dark side of the LDB with its four paws. |
| Open arm entry | Exploratory | The mouse moves from the central platform to an open arm. |
| Closed arm entry | Exploratory | The mouse moves from the central platform to a closed arm. |
| NoGo Head out ** | Risk assessment | While in a protected area, the mouse extends only its head towards the aversive area, then returns to the original position. |
| Go Head out** | Risk assessment | While in a protected area, the mouse extends only its head toward the aversive zone before fully entering it. |
| NoGo Stretch** | Risk assessment | Forward extension of the head and one or both forepaws from a protected area towards an aversive area followed by retraction to the original position. |
| Go Stretch** | Risk assessment | Forward extension of the head and one or both forepaws from a protected area towards an aversive zone before fully entering it. |

*Occurred less than 1% of the time during the LDB and EPM tests.
**In the EPM, head out and stretch postures occur in the closed arms or central platform in direction to the open arm, while in the LDB these behaviors occur when the animal is in the dark compartment and extends its body towards the light side.

**Table S2:** Significant differences in Markov chains' transition probabilities for the Light-Dark Box test. **(A)** GzLM analysis comparing Ctrl and EtOH. **(B)** GzLM analysis comparing Ctrl and Yoh.

### A. EtOH vs. Ctrl

| Transition | | Prob. mean | SD | $X^2_{(1)}$ | p |
|---|---|---|---|---|---|
| Dark entry - HO NoGo | Ctrl | 0.033 | 0.015 | 9.63 | 0.002 |
| | EtOH | 0.020 | 0.012 | | |
| Dark entry - HO Go | Ctrl | 0.011 | 0.012 | 9.76 | 0.002 |
| | EtOH | 0.027 | 0.017 | | |
| Dark entry - ST Go | Ctrl | 0.014 | 0.015 | 8.12 | 0.004 |
| | EtOH | 0.030 | 0.021 | | |
| HO NoGo - HO NoGo | Ctrl | 0.008 | 0.010 | 4.54 | 0.033 |
| | EtOH | 0.002 | 0.004 | | |
| Rearing - walking | Ctrl | 0.138 | 0.036 | 4.12 | 0.042 |
| | EtOH | 0.120 | 0.025 | | |
| ST NoGo - ST NoGo | Ctrl | 0.009 | 0.013 | 5.84 | 0.016 |
| | EtOH | 0.001 | 0.004 | | |

### B. Yoh vs. Ctrl

| Transition | | Prob. mean | SD | $X^2_{(1)}$ | p |
|---|---|---|---|---|---|
| Dark entry - HO NoGo | Ctrl | 0.033 | 0.015 | 7.94 | 0.005 |
| | Yoh | 0.019 | 0.015 | | |
| Dark entry - ST NoGo | Ctrl | 0.024 | 0.015 | 4.57 | 0.032 |
| | Yoh | 0.015 | 0.012 | | |
| Dark entry - ST Go | Ctrl | 0.014 | 0.015 | 6.56 | 0.010 |
| | Yoh | 0.027 | 0.018 | | |
| Sniffing - walking | Ctrl | 0.034 | 0.017 | 5.6 | 0.018 |
| | Yoh | 0.049 | 0.022 | | |
| ST NoGo - HO NoGo | Ctrl | 0.010 | 0.015 | 6.55 | 0.010 |
| | Yoh | 0.001 | 0.004 | | |
| ST NoGo - ST NoGo | Ctrl | 0.009 | 0.013 | 5.47 | 0.019 |
| | Yoh | 0.002 | 0.005 | | |

HO: head out; ST: stretch; Ctrl: control; EtOH: ethanol; Yoh: yohimbine. Ctrl: n = 10/sex; EtOH: n = 10/sex; Yoh: n = 10 females, 9 males.

**Table S3:** Significant differences in Markov chains' transition probabilities for the Elevated Plus Maze test. **(A)** GzLM analysis comparing Ctrl and EtOH. **(B)** GzLM analysis comparing Ctrl and Yoh.

### A. EtOH vs. Ctrl

| Transition | | Prob. mean | SD | $X^2_{(1)}$ | p |
|---|---|---|---|---|---|
| HO NoGo - walking in closed arm | Ctrl | 0.018 | 0.014 | 4.93 | 0.026 |
| | EtOH | 0.007 | 0.012 | | |
| Sniffing in closed arm - rearing | Ctrl | 0.045 | 0.018 | 5.73 | 0.017 |
| | EtOH | 0.033 | 0.013 | | |
| ST NoGo - ST NoGo | Ctrl | 0.007 | 0.009 | 5.7 | 0.017 |
| | EtOH | 0.002 | 0.004 | | |
| Walking in closed arm - HO NoGo | Ctrl | 0.027 | 0.013 | 5.8 | 0.016 |
| | EtOH | 0.015 | 0.014 | | |

### B. Yoh vs. Ctrl

| Transition | | Prob. mean | SD | $X^2_{(1)}$ | p |
|---|---|---|---|---|---|
| Sniffing in closed arm - rearing | Ctrl | 0.045 | 0.018 | 4.23 | 0.04 |
| | Yoh | 0.033 | 0.014 | | |
| ST NoGo - HO NoGo | Ctrl | 0.003 | 0.004 | 6.43 | 0.011 |
| | Yoh | 0.000 | 0.000 | | |
| Walking in closed arm - HO NoGo | Ctrl | 0.027 | 0.013 | 10.36 | 0.001 |
| | Yoh | 0.002 | 0.014 | | |

HO: head out; ST: stretch; Ctrl: control; EtOH: ethanol; Yoh: yohimbine. Ctrl: n = 10/sex; EtOH: n = 10 females, 9 males;  Yoh: n = 10 females, 9 males.

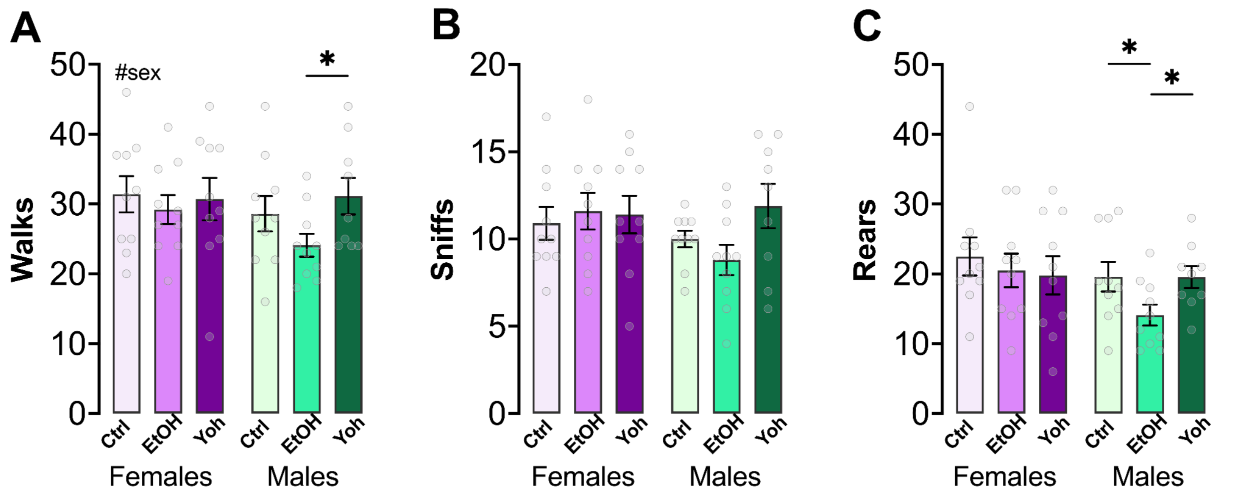


**Figure S1. Exploratory behaviors of adult female and male mice (9th weeks old) in the light side of the LDB five days after saline (Ctrl), ethanol (EtOH) or yohimbine (Yoh) exposures during late adolescence. (A)** The analysis of the number of walks showed a significant effect of sex, as females engaged in more walks than males ($X^2_{(1)} = 4.07$, $p = 0.044$). Sex-separated analyses showed no significant differences among groups in females ($X^2_{(2)} = 0.83$), but males from the EtOH group engaged in less walks than the Yoh group ($X^2_{(2)} = 8.80$, $p = 0.012$; EtOH vs. Yoh: $p = 0.011$). **(B)** There was no significant effect of sex on the number of sniffs ($X^2_{(1)} = 1.99$), and also no group effects following sex-separated analyses (females: $X^2_{(2)} = 0.23$; males: $X^2_{(2)} = 4.44$). **(C)** Rears were increased in females compared to males ($X^2_{(1)} = 9.31$, $p = 0.002$). Sex-separated analyses showed that the number of rears was not affected by treatment in females ($X^2_{(2)} = 1.86$), whereas males in the EtOH group did less rears than Ctrl and Yoh ($X^2_{(2)} = 11.5$, $p = 0.003$; EtOH vs. Ctrl: $p = 0.009$; EtOH vs. Yoh: $p = 0.011$). Ctrl: n = 10/sex; EtOH: n = 10/sex; Yoh: n = 10 females, 9 males.

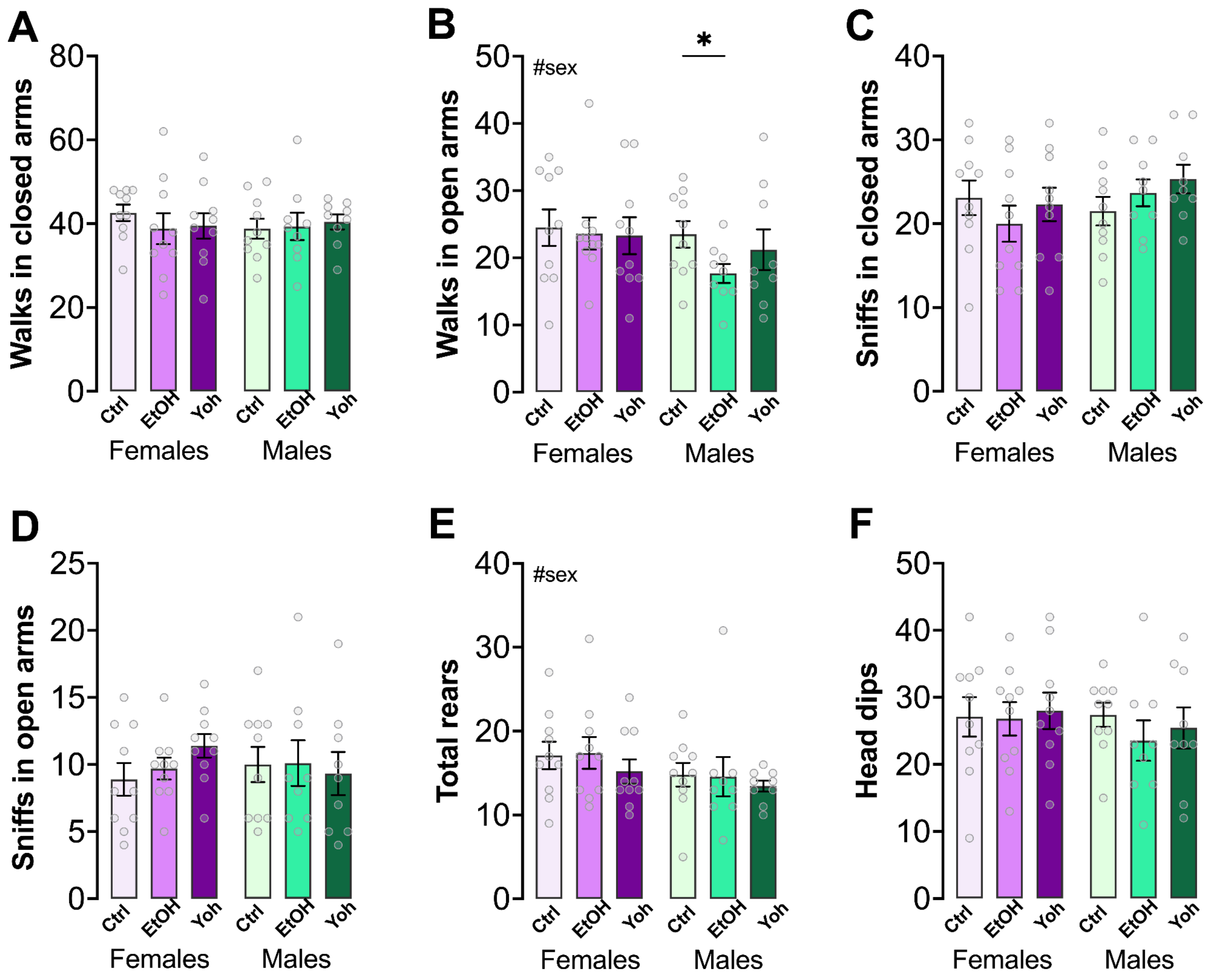


**Figure S2. Exploratory behaviors of adult female and male mice (9 weeks old) in the EPM seven days after saline (Ctrl), ethanol (EtOH) or yohimbine (Yoh) exposures during late adolescence. (A)** Walks in the closed arms were similar in both sexes ($X^2_{(1)}$ = 0.05). Sex-separated analyses showed no group effects (females: $X^2_{(2)}$ = 2.01; males: $X^2_{(2)}$ = 0.33). **(B)** Females walked more in the open arms in comparison to males ($X^2_{(1)}$ = 5.31, p = 0.021), regardless of group. Walks in the open were not affected by treatment in females ($X^2_{(2)}$ = 0.33), whereas EtOH males performed fewer walks in the open arm in comparison to Ctrls ($X^2_{(2)}$ = 7.91, p = 0.019). **(C)** Sniffs in the closed arms did not change across sexes ($X^2_{(1)}$ = 2.61), and sex-separated analyses showed no group effects (females: $X^2_{(2)}$ = 2.40; males: $X^2_{(2)}$ = 3.01). **(D)** Sniffs in the open arms did not change across sexes ($X^2_{(1)}$ = 0.03), and sex-separated analyses showed no group effects (females: $X^2_{(2)}$ = 3.22; males: $X^2_{(2)}$ = 0.33). **(E)** Females did more rears than males ($X^2_{(1)}$ = 4.35, p = 0.037), regardless of group. Sex-separated analyses showed no group effects (females: $X^2_{(2)}$ = 1.74; males: $X^2_{(2)}$ = 0.68). **(F)** Head dips were similar in both sexes ($X^2_{(1)}$ = 1.18). Sex-separated analyses showed no group effects (females: $X^2_{(2)}$ = 0.29; males: $X^2_{(2)}$ = 2.75). Ctrl: n = 10/sex; EtOH: n = 10 females, 9 males; Yoh: n = 10 females, 9 males.

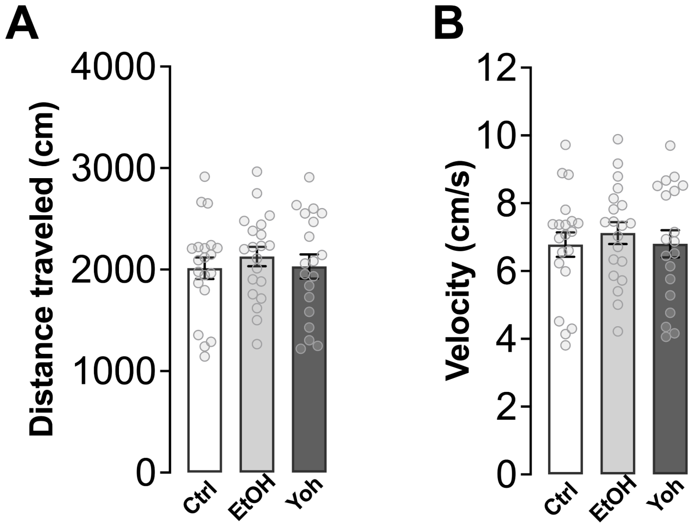


**Figure S3. Distance and velocity during the EPM test following saline (Ctrl), ethanol (EtOH) and yohimbine (Yoh) exposures during late adolescence and adulthood. (A)** Total distance traveled did not differ across groups of mice exposed during late adolescence ($X^2_{(2)}$ = 0.64). **(B)** Velocity was also similar in all groups ($X^2_{(2)}$ = 0.53).
